# Synthetic genetic circuits to uncover and enforce the OCT4 trajectories of successful reprogramming of human fibroblasts

**DOI:** 10.1101/2023.01.25.525529

**Authors:** Katherine Ilia, Nika Shakiba, Trevor Bingham, Ross D. Jones, Michael M. Kaminski, Eliezer Aravera, Simone Bruno, Sebastian Palacios, Ron Weiss, James J. Collins, Domitilla Del Vecchio, Thorsten M. Schlaeger

## Abstract

Reprogramming human fibroblasts to induced pluripotent stem cells (iPSCs) is inefficient, with heterogeneity among transcription factor (TF) trajectories driving divergent cell states. Nevertheless, the impact of TF dynamics on reprogramming efficiency remains uncharted. Here, we identify the successful reprogramming trajectories of the core pluripotency TF, OCT4, and design a genetic controller that enforces such trajectories with high precision. By combining a genetic circuit that generates a wide range of OCT4 trajectories with live-cell imaging, we track OCT4 trajectories with clonal resolution and find that a distinct constant OCT4 trajectory is required for colony formation. We then develop a synthetic genetic circuit that yields a tight OCT4 distribution around the identified trajectory and outperforms in terms of reprogramming efficiency other circuits that less accurately regulate OCT4. Our synthetic biology approach is generalizable for identifying and enforcing TF dynamics for cell fate programming applications.

**One-sentence summary:** Genetic controllers and live-cell imaging offer a versatile strategy for probing the role of transcription factor dynamics in cell fate transitions.

## Introduction

In their landmark papers, Takahashi and Yamanaka demonstrated that murine (1) and human (2) somatic cells can be reprogrammed to a pluripotent state via simultaneous overexpression of OCT4, SOX2, KLF4, and c-MYC, collectively referred to as OSKM. Given the immense potential induced pluripotent stem cells (iPSCs) hold for regenerative medicine and disease modeling, significant efforts have been directed towards developing robust cell fate reprogramming protocols for the efficient and reproducible generation of scalable quantities of high quality iPSCs (3–5). To date, reprogramming of human somatic cells to iPSCs via the ectopic expression of OSKM is still marked by low efficiency (6, 7) and poor quality iPSCs (8– 15), suggesting a knowledge gap regarding how controllable parameters can be optimized to improve the robustness of reprogramming protocols.

Mathematical analysis of the core pluripotency gene regulatory network (GRN) suggests that successful reprogramming hinges on the accurate control of the pluripotency transcription factor (TF) levels (16). The core pluripotency GRN is composed of OCT4, SOX2, and NANOG, three mutually-enhancing and self-activating TFs (17). It has been demonstrated experimentally that fluctuations in OCT4 (18, 19) and SOX2 (20) mediate the destabilization of the pluripotent state. Furthermore, the relative expression levels of OCT4 and SOX2 reportedly influence the efficiency with which somatic cells reach the pluripotent state (21–24). Given that the stability of the pluripotent state depends on key TFs, we hypothesized that the accurate control of total (ectopic and endogenous) TF levels may be necessary to improve the efficiency of reprogramming. This represents a departure from reported reprogramming protocols, which rely on indiscriminate TF overexpression, thereby losing control of TF levels. There are two main drawbacks of these previously published approaches: (a) depending on the gene expression system employed, the level of the exogenously expressed TF can span a very wide range, and (b) once the endogenous genes become reactivated, the total endogenously and exogenously expressed TF level can reach supra-physiological values, thereby overshooting the dosage required for reprogramming. Previous work to control the levels of endogenous TFs relied on either directly perturbing the endogenous genes at the transcriptional level (25, 26) or tuning ectopic expression levels without considering the contribution from endogenous expression (27). An ideal approach would allow for accurate control of endogenous TF levels without interfering with transcriptional remodeling during reprogramming.

Several mechanistic studies have provided in-depth insight into the molecular changes that cells undergo during reprogramming (28–33). These, however, have key limitations: in single-cell RNA sequencing studies (28–32), it is challenging to determine the total expression level of a gene that is expressed both ectopically and endogenously, and RNA levels are not necessarily reflective of protein levels (1, 34, 35); while the quantification of total protein levels is possible with single-cell proteomics sequencing (33), both proteomics and transcriptomics approaches can only be used to reconstruct population-level trajectories as cells must be destroyed for measurements to be taken. In (32), the authors were able to leverage optimal-transport analysis to identify single-cell trajectories; this prior work, however, was done on murine fibroblasts, for which the reprogramming efficiency is higher than for human fibroblasts.

As synthetic biology technologies for controlling gene expression have come of age, there is an opportunity to apply an engineering lens to the problem of reprogramming. Here, we lay the foundation for such an approach by designing two types of genetic circuits, the trajectory generator and the trajectory controller, to reverse- and forward-engineer the expression trajectory of total OCT4, which has been shown to be sufficient for reprogramming of somatic cells to pluripotency and is a central player in the maintenance of pluripotency (21, 36–39). While there have been many works investigating the benefits of epigenetic modifiers, small molecules, OCT4 variants, and the complete omission of OCT4 for enhancing reprogramming efficiency (40–50), here we chose to focus on the OSKM cocktail as a case study to explore the utility of capitalizing on a synthetic biology- and control theory-based approach for improving the efficiency of somatic reprogramming to pluripotency.

For both the trajectory generator and controller circuits, we implemented an miRNA-based strategy that facilitates accurate control over the level of endogenously expressed genes, such as OCT4 (Fig. 1A, left), inspired by the control theory technique of high-gain negative feedback (16, 51). We first designed the trajectory generator, which we delivered to cells via lentiviral vectors such that it generates a wide range of temporal OCT4 dynamics; we tracked these temporal dynamics with clonal resolution over time via high-resolution and high-throughput imaging (Fig. 1A, top right). Our imaging data revealed that a high, stable level of total OCT4 is required for reprogramming success. To validate this finding, we developed the trajectory controller, which precisely fixes the OCT4 dosage at the level of the discovered trajectory during reprogramming (Fig. 1A, bottom right). We found that a trajectory controller that enforces a high OCT4 level yielded the highest reprogramming efficiency when compared to the trajectory generator and trajectory controllers that are not as precise or accurate. The approach presented here is broadly applicable to probing and programming the trajectory of TFs, laying the foundation for future work to improve the robustness of cell fate engineering protocols.

**Figure 1.**
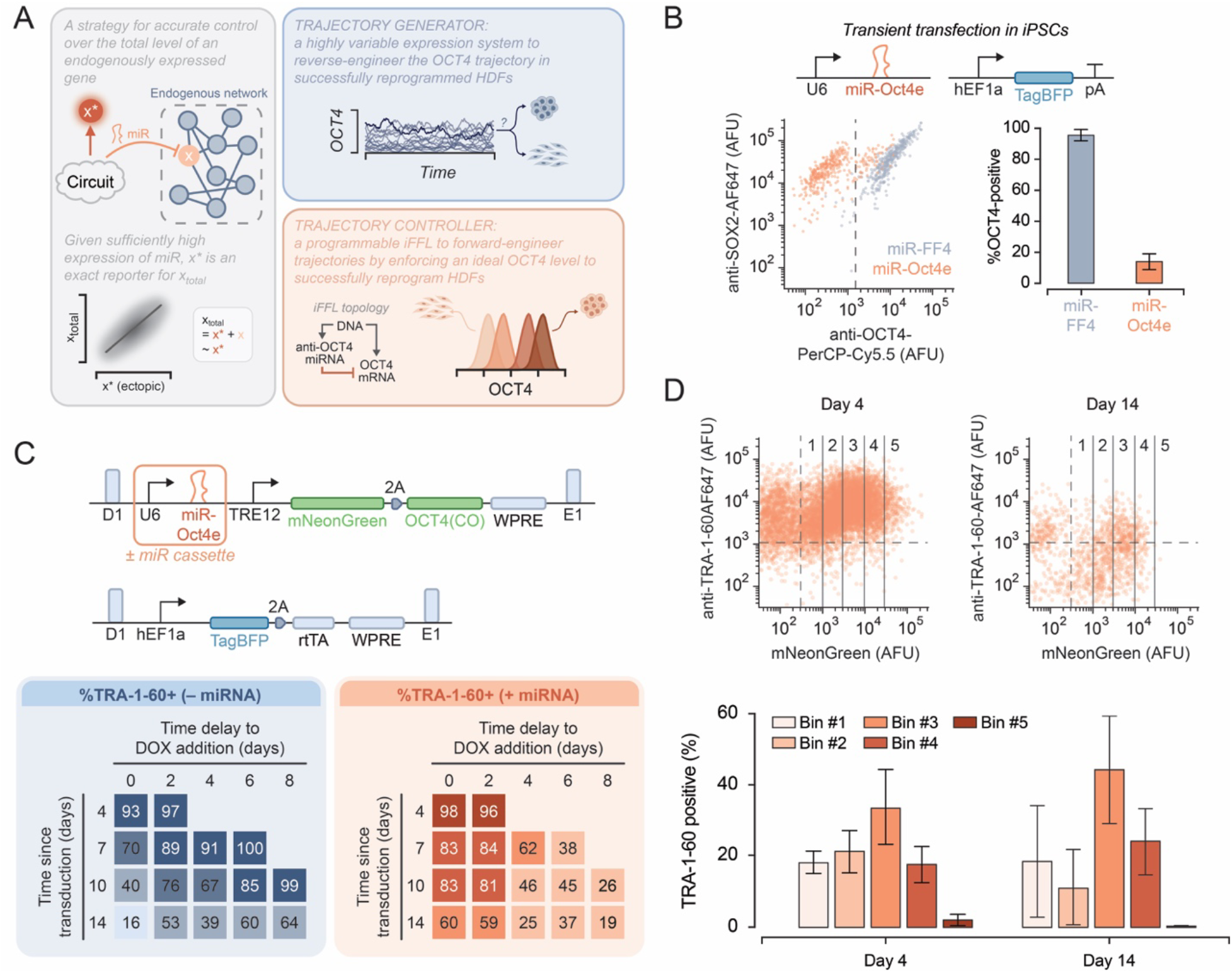
The importance of total OCT4 dosage for the maintenance of pluripotency. **(A)** (left) High-level schematic describing our strategy for implementing a genetic circuit that allows for the accurate control of an endogenously expressed transcription factor (TF). The circuit expresses the ectopic TF (x*) and an miRNA that knocks down the endogenous TF (x) expression. If the miRNA is expressed at a sufficiently high level, then the endogenous contribution x to TF levels is negligible, and the ectopic and total TF (x_total_) levels are approximately equivalent; as a result, we can easily read-out x*, which can be fused to a fluorescent reporter, to reliably determine the level of x_total_. (top right) High-level schematic describing the trajectory generator. (top left) High-level schematic describing the trajectory controller. iFFL: incoherent feedforward loop. **(B)** We transiently transfected iPSCs with the two illustrated plasmids: one encoding U6-driven miR-Oct4e and another constitutively expressing TagBFP, a transfection marker. 48 hr after transfection we stained the cells using PerCP-Cy5.5 anti-OCT4 and AF647 anti-SOX2 antibodies. The scatter plot contains flow cytometry data for transfected (TagBFP-positive) cells. miR-FF4 is a synthetic miRNA that does not target OCT4. The dashed vertical line denotes the OCT4-positive threshold used to calculate the values in the bar graph. Error bars represent the means and standard deviations for n=3 biological replicates. AFU: arbitrary fluorescence units; pA: poly(A) tail. **(C)** We transduced iPSCs with two different versions of the two-lentivirus system shown here, one with and one without the U6-miR-Oct4e cassette, and we cultured them in various doxycycline (DOX) conditions. “Time delay to DOX addition” refers to the number of days post-transduction when DOX was added to the medium. At every timepoint denoted as “time since transduction”, we stained the iPSCs for TRA-1-60. The proportion of TRA-1-60-positive cells for each condition is summarized in the heatmaps. TRE: Tet-responsive element; CO: codon-optimized; WPRE: woodchuck hepatitis virus post-transcriptional regulatory element; rtTA: reversed Tet transactivator (rTetR-NLS-VP64). **(D)** We transduced iPSCs with the version of the system shown in Fig. 1C containing the miRNA cassette and maintained them in DOX-containing medium. We stained the cells for TRA-1-60 four and 14 days post-transduction and analyzed them via flow cytometry. We binned the mNeonGreen population as shown in the scatter plots and determined the proportion of TRA-1-60-positive cells per bin. The horizontal and vertical dashed lines correspond to the TRA-1-60 and mNeonGreen thresholds, respectively. Error bars in the bar graph correspond to the standard deviations of n=3 biological replicates.

## Results

### A strategy for setting total TF levels

In order to probe the role of the dynamics of OCT4 on reprogramming success, we developed a strategy for overwriting endogenous TF levels (Fig. 1A, left). Our system consists of two key features: (a) the ectopic expression of the TF of interest; and (b) the suppression of the endogenous contribution to TF levels with a synthetic microRNA (miRNA) that selectively knocks down the endogenous mRNA species encoding the TF of interest. As this knockdown occurs post-transcriptionally in the cytoplasm, it avoids disrupting transcriptional regulation of the OCT4 gene. To avoid knockdown of the ectopic OCT4 mRNA by the synthetic miRNA targeting the TF coding sequence, we randomized the codons of the ectopic mRNA encoding OCT4.

To implement feature (b), we assayed the knockdown activity of a panel of published (21, 22) and novel miRNAs against OCT4 (Supp. Fig. 1). Our top candidate was the novel miR-Oct4e that, when expressed from the U6 promoter (52), knocked down OCT4 levels by about 24-fold in transfection experiments. To validate that this miRNA could be useful for knocking down physiologically-relevant OCT4 expression, we transfected the U6-driven miR-Oct4e into iPSCs. Indeed, the miRNA eliminated detectable OCT4 expression in over 80% of cells (Fig. 1B).

### Recapitulating the role of OCT4 levels in the maintenance of pluripotency

We next evaluated whether we could leverage this system to experimentally demonstrate the known role that OCT4 levels play in the maintenance of pluripotency (53). To this end, we constructed a two-lentivirus system (Fig. 1C). One lentivirus encodes U6-driven miR-Oct4e and a strong inducible TetR-responsive element (TRE) promoter (Supp. Fig. 2) driving the expression of OCT4 fused to the fluorescent reporter mNeonGreen via a self-cleaving 2A peptide (54). In this system, doxycycline (DOX) activates expression of ectopic OCT4. miR-Oct4e is co-expressed on the same virus as OCT4, such that every transduced cell expresses both components. The second virus contains a constitutive promoter driving the expression of the reversed Tet transactivator (rtTA – rTetR-NLS-VP64) and TagBFP, which serves as a transduction marker. Both lentivirus constructs are flanked by enhancer-blocking insulators (55) to mitigate the extent to which the epigenetic context of lentiviral integration affects transgene expression.

To confirm our ability to perturb the stability of the pluripotent state using this system, we infected iPSCs with two variants of this two-virus system, one with and one without the U6-miR-Oct4e cassette, and tracked the ability of these cells to maintain pluripotency in a variety of DOX conditions. While the miRNA is constitutively expressed, DOX is required for the activation of ectopic OCT4 transcription. To simulate the effects of OCT4 above normal pluripotent levels, we analyzed cells that do not express miR-Oct4e while overexpressing ectopic OCT4. In these samples, induction of OCT4 via DOX addition resulted in a decreased fraction of cells staining positive for the pluripotent stem cell marker TRA-1-60 (11) (Fig. 1C). We also found that the miRNA effectively destabilizes the pluripotent state: in the absence of ectopic OCT4 expression (i.e., in conditions without DOX induction), the iPSCs are unable to compensate for the increased degradation of endogenous OCT4, resulting in the loss of TRA-1-60 staining (Fig. 1C). We then leveraged this experimental system to perform a preliminary investigation of the role of OCT4 dosage on the maintenance of pluripotency. We maintained the iPSCs transduced with the viruses in Fig. 1C in DOX-containing medium, and we found that cells with intermediate-to-high mNeonGreen (a proxy for OCT4) levels are enriched in the TRA-1-60-positive subpopulation (Fig. 1D). Taken together, these results demonstrate that, with our system, we are able to disturb the pluripotency GRN by both increasing and decreasing total OCT4 levels relative to baseline endogenous expression levels and that these perturbations are of sufficient magnitude to trigger the loss of pluripotency.

### A trajectory generator to produce wide OCT4 distributions

These results prompted us to investigate in greater detail how temporal OCT4 dynamics shape the reprogramming process. In particular, we developed a trajectory generator (Fig. 2A) that achieves wide OCT4 distributions and facilitates dynamic tracking of total OCT4 levels over time. As the reprogramming efficiency of human dermal fibroblasts (HDFs) is low, we found that leveraging lineage tracking approaches based on barcoding was not feasible for this application (Supp. Fig. 3). We hypothesized that a live-cell imaging-based approach, which does not require sacrificing cells for measurements and allows us to track the levels of a fluorescent reporter in cells over time, would likely not face the same limitations. This led us to fuse the OCT4 and mNeonGreen proteins to obtain accurate temporal tracking of OCT4 levels (56). In the absence of endogenous OCT4 production, mNeonGreen is an exact reporter for total OCT4 levels. In addition, as OCT4 is a TF, the OCT4-mNeonGreen protein is localized in the nucleus; this simplifies high-throughput image processing and cell tracking as it allows for the spindle-shaped or stellate fibroblasts to be identified by their round nuclei. Previous studies have demonstrated that such a fusion protein does not negatively impact the reprogramming process (57).

**Figure 2.**
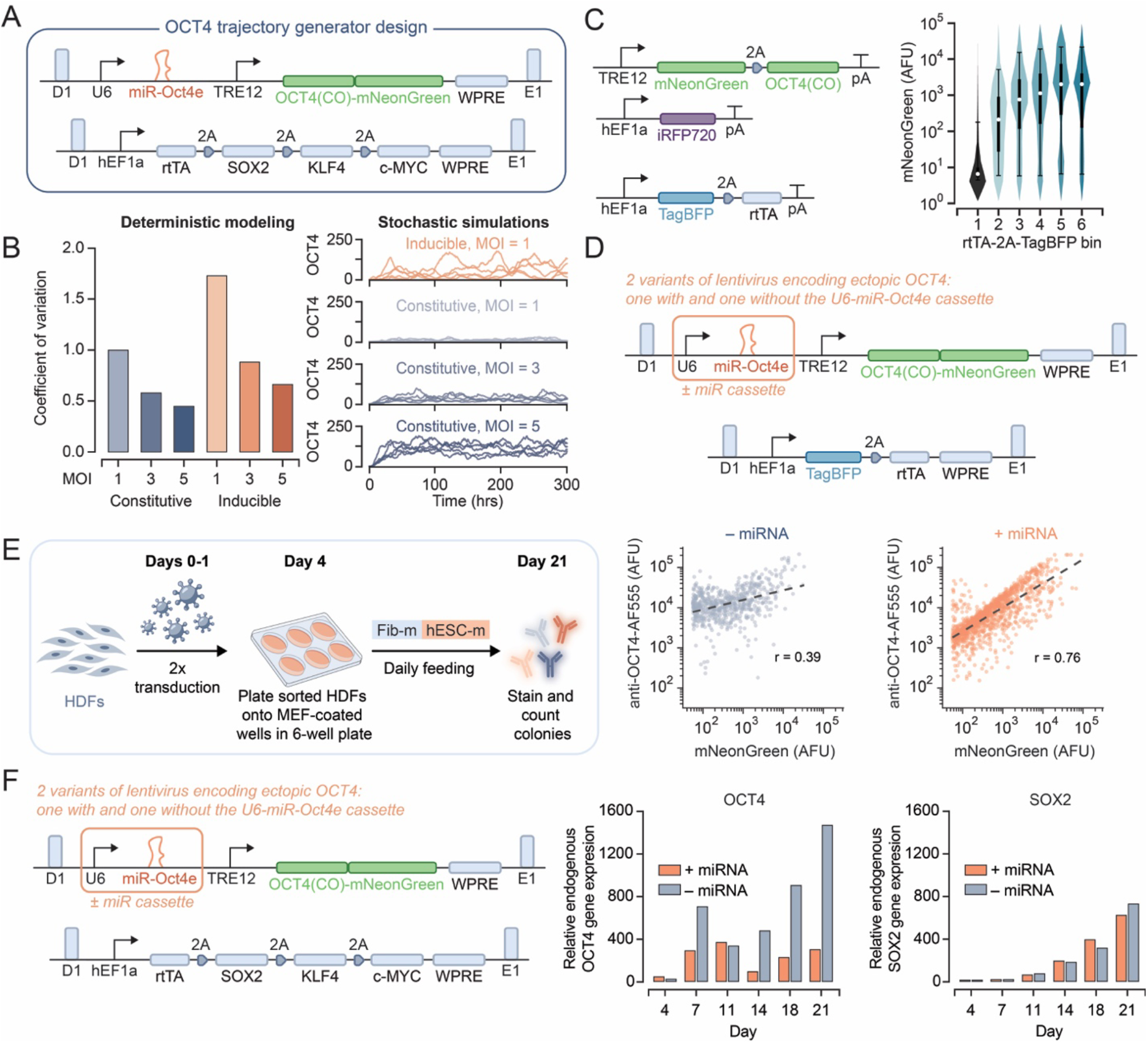
A synthetic genetic circuit for generating variable total OCT4 protein levels during reprogramming. **(A)** The trajectory generator design consists of two viruses: (i) contains a U6 promoter driving expression of miR-Oct4e and an inducible minimal CMV promoter with 12 TetR binding sequences (TRE12) driving expression of mNeonGreen fused to codon-optimized (CO) OCT4, and (ii) contains hEF1a driving expression of rtTA, SOX2, KLF4, and c-MYC. **(B) (left)** We created a deterministic model of the trajectory generator (“inducible”) and an analogous system in which the TRE system is replaced with a constitutive promoter (“constitutive”). We determined the coefficient of variation (CV) of the output OCT4 distribution for three different MOIs for each system. **(right)** We simulated the OCT4 trajectories over time for the same systems at the single-cell level using Gillespie’s Stochastic Simulation Algorithm (SSA) (83). Individual traces represent simulated single-cell trajectories. MOI: multiplicity of infection. **(C)** We performed poly-transfection experiments with two transfection mixes: (i) one containing a plasmid encoding TRE12-driven mNeonGreen and OCT4 linked using a 2A peptide and another plasmid encoding constitutive iRFP720 as a transfection marker; and (ii) a plasmid encoding constitutive TagBFP (transfection marker) and rtTA. Violin plots for a selected low iRFP720 bin illustrate the mNeonGreen output distribution as a function of increasing rtTA dosage. AFU: arbitrary fluorescence units. **(D)** We transduced iPCSs with two versions of the trajectory generator system, one with and one without the U6-miR-Oct4e cassette. We used a modified version of the second virus in Fig. 2A in which TagBFP replaces the SKM factors. We stained the transduced cells for OCT4 four days post-transduction and analyzed them via flow cytometry. miR-Oct4e ensures a high correlation between the reporter mNeonGreen and the TF of interest OCT4. The scatter plots show the double-positive (OCT4, mNeonGreen) cells. The dashed lines represent the linear regression fit of these double-positive points, with the r-value corresponding to each fit shown in the respective plot. AFU: arbitrary fluorescence units. **(E)** Schematic overview of the reprogramming pipeline used for initial characterization of viruses in Fig. 2A. HDFs: human dermal fibroblasts; Fib-m: fibroblast medium; hESC-m: human embryonic stem cell medium. **(F)** We sampled HDFs transduced with the trajectory generator, with and without the miR-Oct4e cassette, at six different timepoints during reprogramming. We conducted RT-qPCR to detect endogenous OCT4 and SOX2, and we calculated OCT4 and SOX2 levels relative to GAPDH. For both reprogramming systems (i.e., with and without the miR-Oct4e cassette), we observed reprogramming efficiency.

We reasoned that, for this type of approach to be viable, we needed a synthetic genetic circuit that promotes gene expression variability. As “burst-like” transcriptional activation is known to lead to wide variance in expression levels (58–61), we reasoned that a switch-like inducible gene expression system, such as with rtTA and DOX, set to intermediate activator expression levels could generate a wide variety of OCT4 trajectories. Via deterministic modeling, we considered the impact of DNA copy number, set by the multiplicity of infection (MOI), on the variability of the OCT4 distribution generated by the inducible trajectory generator in Fig. 2A and an analogous system in which the inducible system is replaced with a constitutive promoter. We found that an inducible system at a low MOI yielded the highest coefficient of variation of OCT4 expression levels in a cell population (Fig. 2B). Stochastic simulations of these circuit topologies confirmed that the inducible system leads to greater temporal variability in OCT4 expression, allowing us to capture a wide range of trajectories (Fig. 2B). To experimentally validate that an inducible system expressed at a low copy number can be used to generate a wide output distribution, we performed poly-transfection experiments, which enable a de-correlation in the delivery of rtTA and rtTA-responsive transcriptional units to cells (62), and found that we can achieve a broader output distribution at low plasmid dosages of each component (Fig. 2C). These results suggest that transducing cells with the inducible trajectory generator system at low MOI would yield highly variable OCT4 levels.

Since mNeonGreen is an exact reporter for total OCT4 levels when endogenously produced OCT4 is strongly knocked down, we next confirmed the ability of low doses of the miR-Oct4e expression cassette to suppress endogenous OCT4 levels. To this end, we transduced iPSCs with the OCT4-containing trajectory generator lentivirus and another lentivirus that constitutively expresses rtTA (without the other reprogramming factors) at low MOIs; we found a strong correlation between mNeonGreen and OCT4 levels only when the OCT4-containing lentivirus also encoded the U6-driven miR-Oct4e (Fig. 2D). These observations demonstrate that, even at low MOI, the U6 and the TRE12 promoters drive sufficiently high expression levels to significantly knock down endogenous OCT4 levels and highly express the transgenic OCT4-mNeonGreen fusion protein, respectively; therefore, we can use the signal intensity of mNeonGreen as an accurate proxy for the total level of OCT4.

We next verified that the trajectory generator is capable of overwriting OCT4 levels during reprogramming by eliminating the endogenous OCT4 contribution throughout a 21-day reprogramming timecourse. To this end, we reprogrammed HDFs according to the pipeline in Fig. 2E with two variants of the trajectory generator, one with and one without the U6-driven miR-Oct4e (Fig. 2F). We sampled the HDFs at various timepoints during reprogramming and measured endogenous OCT4 and SOX2 levels via RT-qPCR. We observed strong activation of endogenous OCT4 in HDFs reprogrammed without miR-Oct4e and found that the miRNA minimizes the endogenous contribution to total OCT4 levels (Fig. 2F). In contrast, the levels of endogenous SOX2, for which we did not implement miRNA-based control, rose during the reprogramming process in both conditions, as expected (Fig. 2F).

### A high-throughput live imaging pipeline to track OCT4 trajectories during reprogramming

We next employed the trajectory generator (Fig. 2A) to reprogram HDFs in a high-throughput imaging setup to track OCT4 levels over time during the reprogramming process (Fig. 3A). We used a CellVoyager7000 confocal imaging system for automated imaging of the transduced HDFs between one and nine times a day. At the end of the timecourse, we stained the cells for SSEA4 and TRA-1-60, surface markers of pluripotent stem cells that have been previously used to gauge colony quality (Fig. 3B,C) (11). As in our previous work, we employed the following classifications of colony quality: Type I colonies are SSEA4^−^, TRA-1-60^−^, Type II colonies are SSEA4^+^, TRA-1-60^−^, and Type III colonies are SSEA4^+^, TRA-1-60^+^ (Fig. 3B,C). Type I and II colonies represent cells in incompletely reprogrammed states while Type III colonies have been shown to be bona fide iPSCs (11).

**Figure 3.**
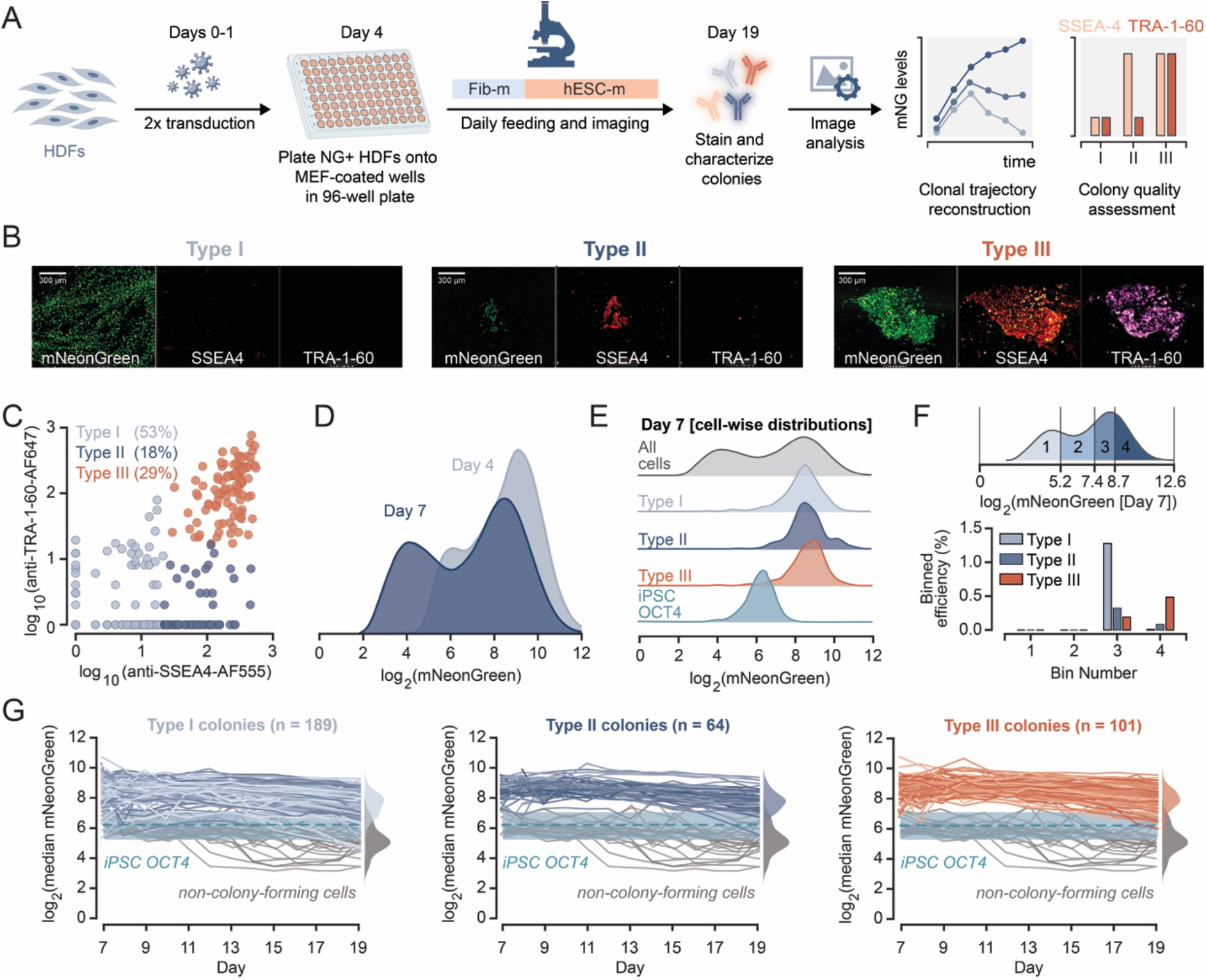
Trajectory reconstruction of reprogrammed HDFs reveals an “ideal” OCT4 trajectory. **(A)** Schematic overview of the high-throughput imaging reprogramming pipeline used for OCT4 trajectory tracking. HDFs: human dermal fibroblasts; Fib-m: fibroblast medium; hESC-m: human embryonic stem cell medium. **(B)** We stained the reprogrammed cells for SSEA4 and TRA-1-60, surface markers that can be used to classify colonies (11). Examples of the staining results are shown in the microscopy images. Scale bar: 300 μm. **(C)** The scatter plot summarizes the frequency of each colony type obtained in the high-throughput imaging reprogramming runs. **(D)** Histograms show the mNeonGreen distribution of the cells infected with the viruses in Fig. 2A four and seven days post-transduction. **(E)** Histograms show the mNeonGreen distribution of the cells infected with the viruses in Fig. 2A at the start of the imaging timecourse. The grey distribution corresponds to all cells present (whether or not they gave rise to a colony). The plotted distributions labeled “Type I,” “Type II,” and “Type III” summarize the cells at the initial time point that gave rise to each of the colony types. The teal distribution corresponds to the OCT4 distribution in iPSCs, as determined by staining both transduced HDFs and iPSCs (see Supp. Fig. 5). **(F)** We binned the initial mNeonGreen distribution into four sections containing equal numbers of cells (top) and calculated the efficiency of reprogramming to Type I, II, and III colonies from cells in each bin (bottom). **(G)** Each line corresponds to the reconstructed trajectory of the median mNeonGreen level for an individual colony. “Non-colony-forming cells” refers to cells that were transduced with the same virus system but did not produce colonies. The dashed line and shaded box correspond to the median plus and minus one standard deviation of the iPSC OCT4 distribution in Fig. 3E. The colored histograms summarize the “colony-wise” distributions, in which each point represents the median OCT4-mNeonGreen expression for a colony corresponding to each type. The gray histograms summarize the “well-wise” distributions for the non-colony-forming cells; for these, each point represents the median OCT4-mNeonGreen expression of all the cells in a well with no colony formation events.

We confirmed with imaging that we could achieve a wide distribution of OCT4-mNeonGreen among transduced cells, and observed a decline in the number of highly expressing cells six days post-transduction compared to four days post-transduction (Fig. 3D), likely due to toxicity associated with high transgene expression levels (Supp. Fig. 4A). Interestingly, all identified colonies, regardless of SSEA4 and TRA-1-60 staining, arose from a narrow subpopulation of relatively highly expressing cells (Fig. 3E). To further probe the effect of OCT4 dosage on reprogramming efficiency, we split the data describing the broad OCT4-mNeonGreen distribution from the first imaging timepoint into four bins, each containing the same number of cells, and calculated the relative efficiency of reprogramming to each type of colony (Fig. 3F). We found that higher levels of OCT4 were enriched in trajectories for all three colony types and, in particular, Type III colonies were produced at the highest efficiency from cells in the highest OCT4 bin.

Next, we validated that mNeonGreen remains an accurate proxy for total OCT4 levels in reprogrammed colonies (Supp. Fig. 5), and we reconstructed the trajectories for each colony present at the last timepoint (Supp. Fig. 6, Supp. Movie 1). Our data suggest that the clonal trajectories for Type I, II, and III colonies are virtually indistinguishable: the majority of trajectories are marked by constant, high OCT4-mNeonGreen throughout reprogramming (Fig. 3G). In contrast, non-colony-forming cells (NCFCs, HDFs that were transduced with the same virus system but did not produce reprogramming-induced colonies) have trajectories that end with low OCT4-mNeonGreen. Taken together, this finding and the observations on the expansion rates of the HDFs infected with the trajectory generator suggest that, in our system, reprogramming HDFs with OCT4 levels that are either too high or too low results in reduced reprogramming efficiency. In particular, when OCT4 levels are too low, the cells are unlikely to reach the pluripotent state (Fig. 3D) and, when OCT4 levels are too high, the cells expand at a markedly lower rate, perhaps due to either cell death or senescence (Supp. Fig. 4A).

### A genetic trajectory controller to enforce precise total OCT4 levels during reprogramming

We next sought to validate that enforcing OCT4 levels precisely within the range of Bin 4 (Fig. 3F) results in higher reprogramming efficiency compared to other OCT4 levels. To this end, we devised a set of four genetic circuit variants that allow for OCT4 expression to be set to different levels in the range covered by our trajectory generator in the imaging runs. We used the trajectory generator as a starting point to develop the trajectory controller (Fig. 4A). While for the trajectory generator we capitalized on the inherent noisiness of an inducible system to generate variability in OCT4-mNeonGreen, here we opted for a constitutive promoter to yield tighter distributions of OCT4-mNeonGreen levels, as predicted by the modeling results in Fig. 2B. To further reduce variation in OCT4-mNeonGreen levels, especially at lower levels of OCT4, we designed an incoherent feedforward loop (iFFL) (63–65) by placing a target site for miR-Oct4e in the 5’ untranslated region (UTR) of the OCT4-containing transcript. We chose to target the 5’UTR rather than the 3’UTR as this placement improves iFFL adaptation to copy number variation (66).

Given that OCT4 levels are controlled by the U6 and constitutive pol II promoters in our system, we reasoned that disparities in the expression dynamics of these two promoters would result in imbalances in the effect of the engineered circuit on total OCT4 levels. To select a constitutive promoter that mitigates this risk, we designed an experimental system to probe the expression dynamics of three constitutive promoters in iPSCs (Fig. 4B). Specifically, we compared the activity of the hEF1a (human elongation factor 1a), cytomegalovirus immediate-early promoter (CMV), and spleen focus-forming virus (SFFV) promoters, as there is evidence of their differential silencing during reprogramming (67–70) or in iPSCs (71). We constructed two sets of lentiviruses: (i) one containing a constitutive pol II promoter (hEF1a, CMV, or SFFV) driving expression of a TagBFP reporter, with or without a U6 promoter driving expression of a synthetic miRNA (miR-FF4), and (ii) one containing a hEF1a-driven mNeonGreen reporter with a target site for miR-FF4 in the 5’UTR. We tracked TagBFP as a reporter for the activity of the pol II promoter and mNeonGreen as a reporter for the activity of the U6 promoter. We transduced iPSCs with these viruses and quantified the expression levels of mNeonGreen and TagBFP over the course of three weeks. We found that the U6 promoter was consistently active in all the experimental conditions, and that during this time, we observed silencing of CMV and SFFV, but not of hEF1a (Fig. 4B). Since we found that neither the U6 nor the hEF1a promoters silenced during this experiment, we selected the hEF1a promoter for our trajectory controller design.

**Figure 4.**
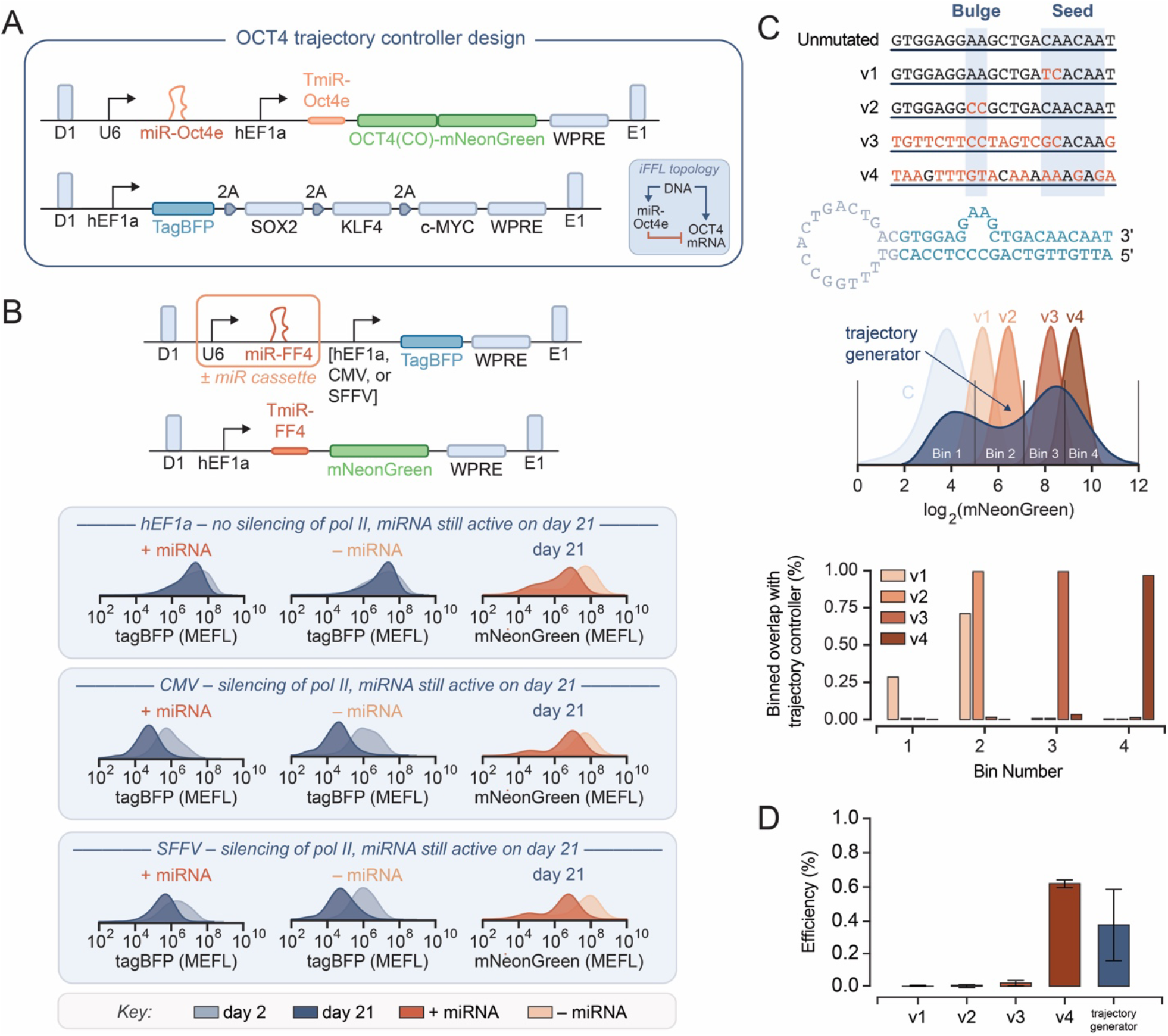
Enforcing the “ideal” OCT4 trajectory via an OCT4 trajectory controller validates role of OCT4 dosage on reprogramming efficiency. **(A)** Schematic of the trajectory controller design. TmiR: miRNA target site. **(B)** We transduced iPSCs with six versions of the illustrated two-virus system: containing one of the constitutive promoters listed (hEF1a, CMV, or SFFV) driving TagBFP expression; in these viruses, the U6 cassette was either included (+ miRNA) or omitted (– miRNA). We analyzed the transduced cells via flow cytometry two and 21 days post-transduction. MEFL: molecules of equivalent fluorescein. **(C) (top)** Sequences of miR-Oct4e target site variants with point mutations (with respect to the unmutated control, a perfectly complementary target site) indicated in orange text. The mutants were generated by using the following nucleotide substitutions: *A* ↔ *C, T* ↔ *G*. The ‘bulge’ and ‘seed’ regions denote sequences that are critical for miRNA-mRNA binding (84). **(middle)** The histograms summarize the OCT4-mNeonGreen distributions of HDFs infected with variants of the virus system in Fig. 4A, seven days post-transduction. “Trajectory generator” refers to the two-lentivirus system in Fig. 2A. “C” is the mNeonGreen distribution of untransduced HDFs (i.e., autofluorescence). The vertical lines demarcate the same bins used in the analysis in Fig. 3F. **(bottom)** For each variant, we determined the proportion of the mNeonGreen distribution that overlaps with each bin from Fig. 3F. **(D)** We reprogrammed HDFs using the trajectory controller variants characterized in Fig. 4C. We stained cells for TRA-1-60 at the endpoint and determined the number of TRA-1-60-positive (Type III (11)) colonies. We calculated the reprogramming efficiency as the ratio of TRA-1-60-positive colonies and the number of cells plated per well on day 4. The values in the bar graph correspond to the mean efficiencies, and the error bars represent the standard deviations for n=3 biological replicates.

To validate our finding that high OCT4 levels within the ranges of Bin 4 are more conducive to reprogramming (Fig. 3F,G), we created four trajectory controller variants (v1, v2, v3, and v4) with increasingly mutated miR-Oct4e target sites in the 5’UTR of OCT4-mNeonGreen (Fig. 4C, top). As the knockdown strength of miRNAs is dependent on complementarity of the target mRNA sequence to key segments of the miRNA (72), target sites with partial complementarity can be employed to decrease the repression strength and thereby precisely set the expression levels of a gene of interest (27). To capture a range of OCT4 levels with these trajectory controllers comparable to that of the trajectory generator, we used a high MOI for the OCT4-containing lentivirus; it is known that an increase in the MOI also has the benefit of reducing the width of the output distribution, as confirmed by our modeling results (Fig. 2B). Despite this increase in the MOI, the HDFs infected with each variant expanded comparably to HDFs transduced with the trajectory generator for a given OCT4 expression level (Supp. Fig. 4B), consistent with previous reports of the relationship between reprogramming factor dosage and HDF expansion (73). We transduced the HDFs with the SKM-containing virus at the same low MOI for the experiments using the trajectory generator (Fig. 3) and the trajectory controllers (Fig. 4) for consistency and determined that SOX2 expression levels were comparable across the trajectory generator and controller reprogramming systems (Supp. Fig. 7).

The distribution of OCT4-mNeonGreen levels from the cells transduced with the trajectory controller variants overlap in the intermediate-to-high range of expression of the cells infected with the trajectory generator (Fig. 4C, middle). Applying the bins from Fig. 3F to these distributions, we found that the trajectory controller variant v4 generates a tight distribution that overlaps by 97% with Bin 4 (Fig. 4C, bottom), from which the trajectory generator had yielded the highest fraction of Type III colonies. Based on this, we hypothesized that the v4 controller would provide the highest reprogramming efficiency compared to the imprecise trajectory generator and the other precise but not accurate trajectory controller variants. To confirm this, we performed a bulk reprogramming experiment using the protocol outlined in Fig. 2E for each of the trajectory controller variants and stained for TRA-1-60 to determine the number of Type III colonies (11). Indeed, the v4 system reprogrammed HDFs to iPSCs at higher efficiency than the other variants as well as the trajectory generator, with a 29-fold increase in efficiency compared to the second-best performing trajectory controller variant, which generates a tight OCT4 distribution but with a slightly lower median compared to v4, and a 1.7-fold increase in efficiency compared to the trajectory, which generates a very wide OCT4 distribution and overlaps with the v4 distribution (Fig. 4D). Overall, our results demonstrate that by identifying ideal trajectories for TF expression within a given reprogramming context and building genetic circuits that precisely enforce these trajectories, we can more efficiently drive desired cell fate conversions.

## Discussion

Here, we sought to explore whether accurate control over TF levels can drive more efficient cell fate programming outcomes. Reprogramming of HDFs to iPSCs served as a testbed for us to implement genetic controllers that allow exploring and enforcing TF trajectories associated with successful cell fate transitions. The application of these tools to hiPSC reprogramming, a well-studied yet inefficient process with potential clinical applications, revealed tight control of OCT4 about an ideal level as a requirement for higher reprogramming efficiency.

To this end, we devised genetic controllers that provide total control over the OCT4 level, including the ability to tune both expression variability and average levels. Our controller designs enable the overwriting of the levels of endogenously expressed TFs via a miRNA-based strategy (Fig. 1). We devised two sets of genetic circuits to achieve two distinct goals: (a) an inherently noisy DOX-inducible system to generate a variety of total OCT4 trajectories and evaluate the effect of temporal OCT4 dynamics on reprogramming efficiency (the trajectory generator, Fig. 2), and (b) an miRNA-based iFFL that facilitates accurate tuning of OCT4 levels in order to improve reprogramming efficiency (the trajectory controller, Fig. 4). Using these synthetic genetic circuits, we demonstrated that by reverse- and forward-engineering ((Fig. 3) and (Fig. 4), respectively) the trajectories of successful reprogramming clones, we can more efficiently control the transition to pluripotency.

We envision that our high-throughput live-cell imaging pipeline and genetic circuit designs can be leveraged to study the roles of the temporal dynamics of key TFs and improve the efficiency of other cell fate conversion applications, such as differentiation and transdifferentiation. This is because the circuit architectures described herein can be easily extended to control any endogenous TF of interest. Our approach also provides the first control theory-based strategy (16) to accurately steer the levels of key lineage-specifying TFs to desired values. Although we focused solely on the OSKM cocktail for iPSC reprogramming as a case study, our approach can be used synergistically with other reprogramming methods that rely on different combinations of TFs, leverage small molecules and culture conditions, and incorporate epigenetic modifiers to further improve efficiency (40–50). Another natural extension of this work entails multiplexing our genetic controllers for simultaneous control of multiple endogenous genes with different sets of miRNAs. Our high-throughput imaging approach could be leveraged as the starting point for this type of future work. In doing so, we could gain a deeper understanding into how levels and dynamics of key TFs influence reprogramming or other cell fate conversions. We expect that our strategy will have broad utility both in regenerative medicine and cell therapy, where better-controlled expression of endogenous TFs of interest could be leveraged to improve the production, safety, and efficacy of emerging therapies (74).

## Acknowledgements

We are grateful to Dr. Jeff Wrana for generously sharing his lineage tracking barcode library. We also thank the flow cytometry core at the Koch Institute’s Robert A. Swanson (1969) Biotechnology Center for technical support. We thank Dr. Raphaël V. Gayet for helpful comments on the manuscript.

## Author contributions

Conceptualization: KI, NS, TB, RDJ, JJC, RW, DDV, TMS; methodology: KI, NS, TB, RDJ, MMS, EA, SB, JJC, RW, DDV, TMS; software: KI, NS, TB, TDJ, SB; validation: KI, NS, TB, RDJ, TMS; formal analysis: KI, NS, TB, TDJ, TMS; investigation: KI, NS, TB, TDJ, TMS; resources: JJC, RW, DDV, TMS; data curation: KI, NS, TB, RDJ, TMS; writing – original draft: KI; writing – review & editing: KI, NS, TB, RDJ, SB, JJC, RW, DDV, TMS; visualization: KI, TMS; supervision: JJC, RW, DDV, TMS; project administration: JJC, RW, DDV, TMS; funding acquisition: JJC, RW, DDV, TMS.

## Data and materials availability

All data needed to evaluate the conclusions in the study can be found in the paper and the supplementary materials. Plasmids will be made available to the scientific community through Addgene. Correspondence and requests for materials should be addressed to TMS or DDV.

